# Northern palsa and peat ecosystem microbiomes are potentially supported by hydrogen and carbon monoxide metabolism

**DOI:** 10.64898/2026.11.09.765253

**Authors:** Nea Tuomela, David N. Thomas, Hermanni Kaartokallio, Tommi Välikangas, Jenni Hultman

## Abstract

Northern peatland and permafrost ecosystems are facing rapid changes, with increasing temperatures, permafrost thaw and shifts in hydrology. Characterizing microbial communities and metabolic potential from the soils and associated water bodies is crucial to understanding the changes in biogeochemical cycling in these ecosystems. For this, samples were collected and sequenced from subarctic palsa and peat soils, along with samples from associated thaw ponds and streams. Metagenomics was utilized in describing the microbiome and its metabolic potential in these environments. Palsa soils hosted microbial communities that were distinct from the peatland soils, ponds and stream waters. Soils and their adjacent ponds shared taxa, and peat soils and ponds also shared taxa with the streams, unlike palsa soils which were likely more hydrologically isolated. Based on metabolic marker gene ratio for denitrification [(*nirS*+*nirK*)/*nosZ]*, palsa soils had the strongest potential to be a source of nitrous oxide (N_2_O) out of all the sample types. Marker genes for hydrogen gas (H_2_) and carbon monoxide (CO) utilization were abundant across soil environments and metagenome assembled genomes, suggesting that trace gas oxidation is likely one of the key energy sources for microbiomes. In the palsa ponds and peat soils *pmoA*, marker gene for methanotrophy, was the most abundant compared to the other sample types.

## Introduction

Northern peatlands and permafrost are the largest reservoirs of terrestrial organic carbon and nitrogen, and they are facing drastic changes as the Arctic warms rapidly (Hugelius et al., 2020; Schuur et al., 2022; Treat et al., 2024). Permafrost and peat soil stabilize carbon and nitrogen with cold temperatures or frozen states preventing or slowing down degradation. However, as permafrost thaws the organic matter becomes available to microbes (Hultman et al., 2015). Microbes are central in many of the elemental cycles, including carbon and nitrogen and microbial processes are largely responsible for the production and consumption of greenhouse gas emissions such as CO_2_, CH_4_ and N_2_O (Ernakovich et al., 2022; Hultman et al., 2015).

In subarctic regions, permafrost formations can consist of isolated permafrost, such as palsas. These are permafrost mounds or plateaus, where a permafrost layer is insulated by an overlaying dry peat (Seppälä, 2011). Palsa mires and other isolated permafrost sites are highly vulnerable to climate change, and they are rapidly degrading across the arctic and subarctic regions (Borge et al., 2017; Luoto & Seppälä, 2003; Mamet et al., 2017): e.g. An areal decrease of up to 71% of the palsa mires in Norway between the 1950s and 2010s has been measured (Borge et al., 2017). Thaw ponds and thermokarst fen are formed following palsa degradation. The thaw ponds are considered to be hotspots of greenhouse gas emissions as they receive high loading of organic matter from the thawing soils (Chauhan et al., 2024; Knutson et al., 2025; Roiha et al., 2015; Tuomela et al., 2025).

Permafrost and peat soils host a diverse community of bacteria and archaea, including methanogens, methanotrophs, nitrifiers and denitrifies (Waldrop et al., 2025). In intact permafrost, microbes are expected to be in static state and low activity levels, due to cold conditions and low availability of nutrients (Waldrop et al., 2025). But as permafrost thaws, the active microbial communities shift rapidly within days with increased abundances of groups such as denitrifiers, methanotrophs and methanogens (MacKelprang et al., 2011).

Trace gas metabolism, namely metabolism of H_2_, CO and CH_4_, has been detected widely across different soils, including peatlands (Välikangas et al., 2026), tundra (Viitamäki et al., 2022), Antarctic soils (Lebre et al., 2023; Ray et al., 2022) and volcanic soils (Fantom et al., 2025). Trace gas oxidizers enhance the biodiversity and the resilience of microbial communities. By utilizing both trace gases and organic energy sources, microbes are granted an advantage as they can flexibly utilize both energy sources and switch from organic compounds to trace gases (Bay et al., 2021; Greening & Grinter, 2022). Utilizing trace gases can also increase the carbon use efficiency of bacteria (Greening & Grinter, 2022). While described in many ecosystems, to our knowledge the marker genes for trace gas utilization have not been described in sporadic permafrost environments.

While microbiomes of permafrost and the emerging thaw ponds have received increasing attention, these two highly interlinked environments are generally studied separately (Chauhan et al., 2024; Waldrop et al., 2025; Woodcroft et al., 2018; Wurzbacher et al., 2017). Studies with permafrost thaw sequence generally include the transect from permafrost soil to thermokarst peat (Emerson et al., 2018; Hultman et al., 2015). Studies with the transect of terrestrial and aquatic habitats remain scarce (Burnett et al., 2026; Crump et al., 2012; Leroy et al., 2025, 2026), yet they are highly pertinent to in understanding the carbon exchange of the whole peatland (Kuhn et al., 2018).

In this study, metagenomics was utilized to characterize the microbial community composition and metabolic potential in a subarctic, sporadic permafrost peatland environment, including soil and water samples. Our aim was to describe the community composition in the continuum from soil to highly connected ponds and streams, in both permafrosts affected palsa and an adjacent peatland with no permafrost. We hypothesize that the palsa and peatland environments differ in both microbial community composition, but that the soil and adjacent waters share taxa. We also hypothesize that the potential for trace gas oxidation is widespread across all the sample types.

## Methods

### Sampling site and soil chemical properties

We sampled two palsa mounds and peatland soils in Vaisjäeggi palsa mire and peatland in Ohcejohka (Utsjoki) Finland during June 2023, which was in conjunction with the end of the spring snowmelt. The palsa mire consists of isolated degrading palsa mounds surrounded by thaw ponds. The peatland site has a shallow peat layer (12cm) over mineral layer. The site is described in more detail by Tuomela et al. (2025).

Six replicate soil profiles were collected from two palsa mires by hammering a steel corer into the soil up to 56-81cm. Soil cores were sliced into 2-5cm slices (n=72) with an ethanol sterilized knife. The soil profiles were divided into active and frozen, based on the depth where the frozen soil began (depth 21-33cm). Peat soils were sampled from 2, 4 and 10cm depths (n=15). The samples were stored in sterile plastic bags on ice until they were transferred to -80 °C freezer within a few hours.

Water samples were collected, stored on ice in dark and filtered within few hours through 0.45 and 0.1 µm pore size and the filters stored at -80 °C (Tuomela et al., 2025). The sample types were divided into following sample types: palsa active (n=20), palsa frozen (n=72), palsa pond (n=6), peat soil (n=15), peat pond (n=5) and stream water (n=10).

Soil physicochemical properties were measured from lyophilized and ground soils. Soil pH was measured according to the international standard ISO10390. Loss on ignition (LOI) was used to analyze soil organic matter (SOM) content according to Finnish standard SFS3008. Soil total carbon and total nitrogen analysis was performed with a high temperature induction furnace (LECO, 828 series). The differences in physicochemical properties between sample types were tested with Kruskal Wallis test followed by Dunn’s test with Bonferroni corrections.

### DNA extraction and sequencing

DNA extraction was performed for 0.25-0.30 g of soil from the inner part of the sliced core, avoiding the surface of the sample, with DNeasy PowerSoil Pro Kit (Qiagen, Hilden, Germany). Cell disruption step was done with FastPrep homogenizer (MP Biomedicals, Heidelberg, Germany; 30s, 5.5m/s). The rest of the extraction was completed following the manufacturer’s instructions. For water samples, the same protocol as described by Tuomela et al., (2025) was followed for the sequential filters, which were combined. Libraries were prepared with Nextera™ DNA Flex Library Preparation Kit (Illumina, San Diego, CA, USA) and paired-end sequencing was done with AVITI (Element Biosciences, 2x150bp) at the Institute of Biotechnology, University of Helsinki.

### Bioinformatical analysis

#### Quality control and read-based analysis

Raw sequences were trimmed using cutadapt v4.9 to remove Nextera adapters. Quality was checked with FastQC v0.11.9 and MultiQC v1.19. After trimming, the sequences from individual lanes were combined for each sample. For taxonomic annotation, trimmed and combined reads were mapped against GTDB-tk r232 with sylph v.0.9.0 (Shaw & Yu, 2024). The R-package vegan 2.7-6 (Oksanen et al., 2026) was used to ordinate the relative abundance of bacteria and archaea species, with Nonmetric Multidimensional Scaling (NMDS) and visualize the resulting ordination.

To investigate the metabolic functional potential, the metagenomic reads were annotated using the curated 51 metabolic marker gene database from Greening Lab (Leung and Greening 2021, Lappan et al. 2023) similar to (Lappan et al., 2023; Välikangas et al., 2026). Briefly, the paired-end reads were aligned against the compiled metabolic marker gene database with DIAMOND (Buchfink et al., 2014) blastx and filtered according to the marker gene specific minimum percentage identity thresholds following (Lappan et al. 2023). The filtered paired-end alignment hits were reconciled at the paired-fragment level so that each sequenced fragment contributed at most to one marker gene assignment, transformed into gene length- and sequencing depth normalized abundance measure (length adjusted reads per million (L-RPM)) and aggregated by marker gene. For more details, see Supplementary Methods.

#### Metagenome assembled genomes

The metagenomes were assembled into 9 coassemblies with MEGAHIT v1.2.9 (min-contig-length 1000, k-min 57, k-max 157, k-step 10) (Li et al., 2015). Coassemblies were mainly compiled by sample types. The coassembly quality was checked with MetaQuast v.5.2.0 (Gurevich et al., 2013). The assemblies were then processed following anvio metagenomics workflow with anvio v8 (Eren et al., 2020), with contigs ≥ 1500bp retained. Reads were mapped back to assemblies with Bowtie2 v.2.5.3 (Langmead & Salzberg, 2012) and indexed with SAMtools v.1.21 (Li et al., 2009). Sample profiles were generated (anvi-profile) and merged (anvi-merge). Manual binning was done with anvi-cluster-contigs with MetaBAT2 v.2.15 (Kang et al., 2019) and then manually refined. MAGs were dereplicated with fastANI v.1.34 (similarity threshold 0.98) (Jain et al., 2018) and quality checked with CheckM2 v.1.1.0 (Chklovski et al., 2023), following MIMAG standards of completeness ≥ 50 % and contamination <10 % (Bowers et al., 2017). Taxonomy was assigned with GTDBtk v. 2.7.2 (database v.r232) (Chaumeil et al., 2020) using the classify workflow.

CoverM v.0.7.0 (Aroney et al., 2025) was used to determine abundance of the MAGs across samples in transcripts per million (TPM) (methods tpm covered_fraction). Only MAGs with a covered fraction ≥ 25% were considered as present. Unique genera between the sample types were visualized with UpSet plot using R-package ComplexUpset v1.3.3. (Lex et al., 2014). The community dissimilarity based on MAG level abundance was explored with NMDS using Bray-Curtis dissimilarities and the resulting ordination was visualized.

Metabolic potential within the MAGs was explored by annotating the MAGs with anvi’o v9 (Eren et al., 2020) programs anvi-run-kegg-kofams and anvi-estimate-metabolism against KEGG v.116. Metabolic pathways and modules related to methane and nitrogen metabolism were inspected, and the metabolic module or pathway was considered as complete if pathwise completeness was over 75%.

Metabolic marker genes were also identified from the predicted genes of the recovered final MAGs using the same curated compiled metabolic marker gene database as in the read-based analysis (Leung and Greening 2021, Lappan et al. 2023). Predicted nucleotide sequences from genes in each MAG were aligned against the metabolic marker gene database protein sequences using DIAMOND (Buchfink et al., 2014) blastx v2.2.5.185 (--max-target-seqs 1 --max-hsps 1 --outfmt 6 qseqid stitle pident length qstart qend sstart send evalue bitscore qcovhsp scovhsp slen) similar as in (Lappan et al., 2023). The raw alignment hits were further filtered following the predicted protein approach of (Lappan et al., 2023). Alignments were retained when either query or subject coverage was at least 80%, followed by marker gene specific minimum percentage identity filtering. The percentage identity thresholds were the same as those used for predicted proteins by (Lappan et al., 2023): identical thresholds as used in the read-based analysis except for *AtpA* and *PsbA* (60%), *RdhA* (45%), *Cyc2* (35%), and *RHO* (30%) and NiFe Group 4 hydrogenases were required to have at least 60% percentage identity. These filtered MAG marker gene alignments were subsequently used to summarize the presence the metabolic marker genes among the recovered final MAGs.

## Results

### Physicochemical properties differed between palsa and peat soils

The palsa soils had pH’s of 3.80±0.09 (mean and standard deviation) and 4.06 ±0.47 in active and frozen layers, respectively (Figure 1). The palsa profile 26 had an increase in pH from 4 to 5 in the lowermost sampled depth of 40-80 cm (Table S1). The soil organic matter content (SOM) was high in palsa soils, 97.3±2.78% in active and 97.9 ±0.69% in frozen layers. Total carbon to total nitrogen (C:N ratio) was 38.7 ±5.67 in active and 36.1±6.49 in frozen layers. Palsa active and frozen soil were not significantly different from each other in pH, SOMor their total carbon to total nitrogen ratio (C:N ratio). Palsa soil samples had similar environmental properties as in literature (Kirkwood et al., 2021; Voigt et al., 2017). However, the palsa soils in this study had slightly lower pHs than palsa soils described in Hudson Bay (pH 4.8, Kirkwood et al., 2021).

**Figure 1.**
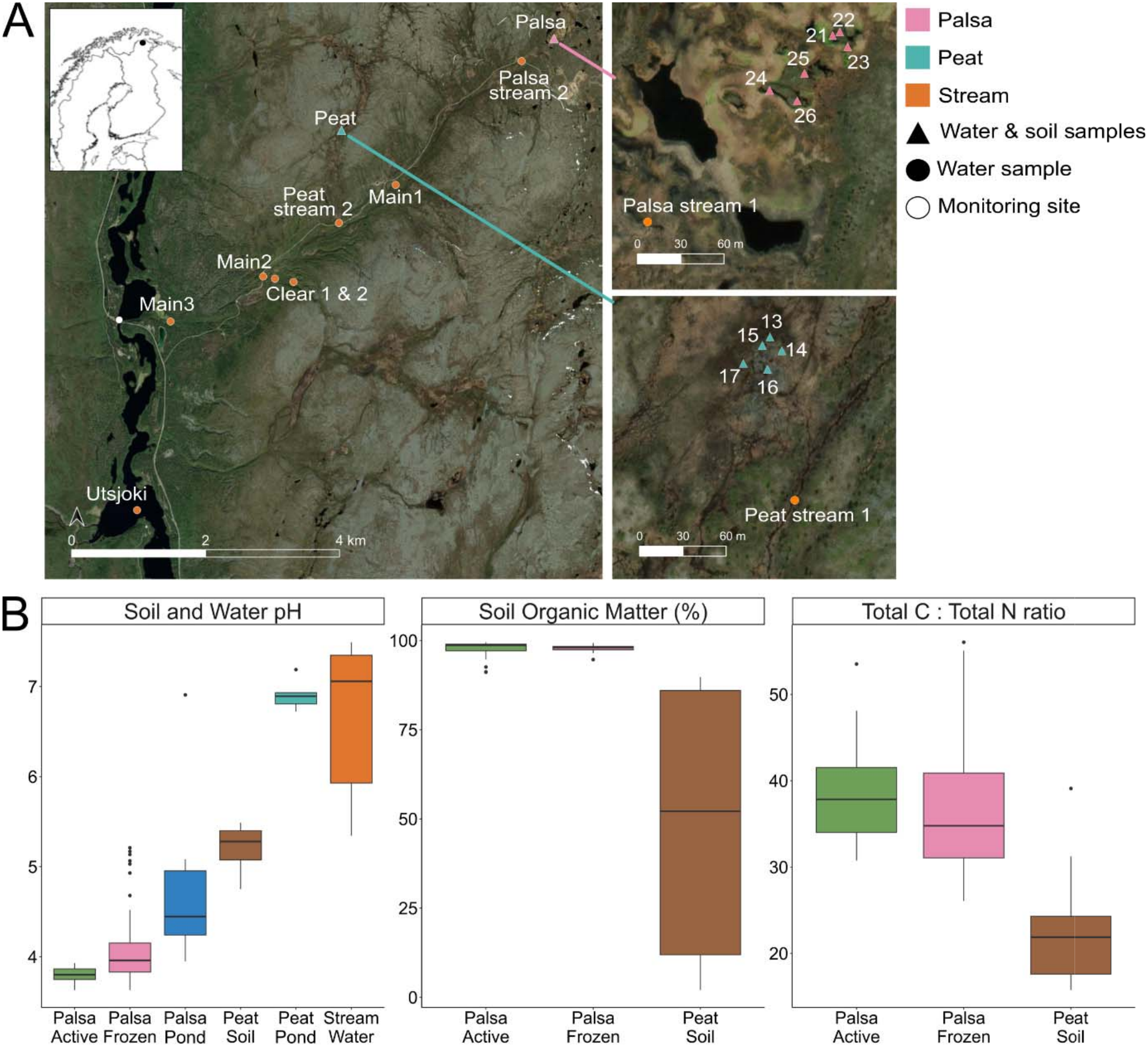
Map of the sampling sites and long-term monitoring site Skállovárri in Kevo, Utsjoki region in Finland (A) and the measured properties of soil and water samples including pH, soil organic matter (SOM) and total carbon to total nitrogen ratio (B).

In the peat soils, pH was 5.22 ±0.23 (mean and standard deviation), SOM was 48.57 ±36.73% and the C:N ratio was 22.39 ±6.42. Both active and frozen palsa layers had significantly higher SOM and C:N ratio than peat soils and both palsa soil types were significantly more acidic than any of the sample types, including the palsa waters (Dunn’s test, p<0.05). Palsa ponds had an average pH of 4.8, while in peat ponds and associated streams the pH was 6.9 and 6.7 respectively (Tuomela et al., 2025).

### Microbial community composition differs in palsa soils and ponds compared to the peatland ponds, soils and stream waters

Ordination of the read-based taxonomy showed that the sample type explained much of the variation among the samples (Figure 2). Active and frozen palsa soils had distinctive community compositions compared to those from peat soils and water samples. Notably, the deepest palsa soil layers (profile 26, depth >40cm) were separating from the frozen palsa soils and were more similar to the palsa waters. These soil layers were dominated by *Halobacterota* and *Caldicericota* (Figure S1). In other palsa soil samples and peat soils, the communities were dominated by *Acidobacteriota, Actinomycetota, Pseudomonadota* and *Chloroflexota*. *Acidobacteriota*, *Pseudomonadota*, *Patescibacteriota*, *Desulfobacterota* and *Bacteroidota* were the most abundant phyla in water samples (Figure S1).

**Figure 2.**
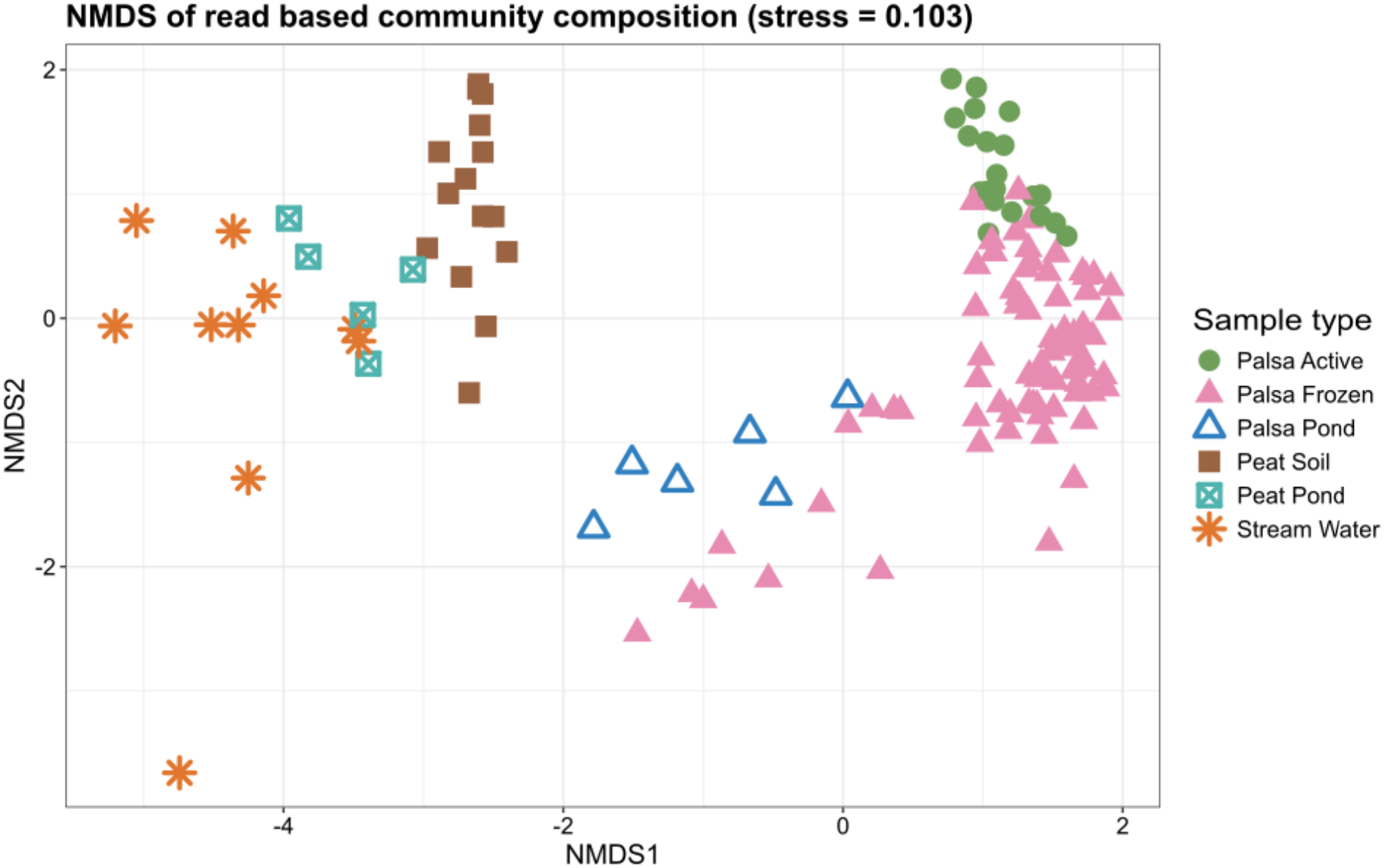
Non-metric Multidimensional Scaling (NMDS) with Bray Curtis distance of the microbial community composition of the investigated northern palsa soil, peat soil, palsa pond, peat pond and stream samples. The ordination was based on Bray-Curtis dissimilarities of species-level relative abundances of read-based taxonomy data.

### Metabolic marker genes suggest incomplete denitrification potential in palsa soils and trace gas metabolism supporting microbial growth across soil types

#### Nitrogen metabolism

Marker genes related to denitrification and N_2_O production (*nirK*, *nirS*, *norB*) were identified in the palsa frozen soils, peat soils and stream waters (Figure 3). The N_2_O reduction marker gene (*nosZ*) was detected in low abundance in the palsa soils, except for palsa soil profile 26 deeper layers. Overall peat soils had the highest L-RPM values of *nosZ.* The ratio of (*nirS*+*nirK)/nosZ*) indicates the potential for N_2_O production over N_2_O retention (Saarenheimo et al., 2015). The ratio was consistently >1 in all samples, with highest in the palsa active layer soils (mean and standard deviation 15.9 ±12.0 L-RPM) and frozen palsa soils (6.1 ±3.5 L-RPM), followed by the stream waters (4.2 ±1.1 L-RPM), the peat ponds (4.2 ±1.2 L-RPM) and lowest in the palsa ponds (3.7 ±1.9 L-RPM). The peat soil had the lowest ratio (3.1 ±1.3 L-RPM). Notably within the frozen palsa soil samples, the deepest layers of palsa soil profile 26 had a higher abundance of *nosZ* than the other palsa or peat soil samples.

**Figure 3.**
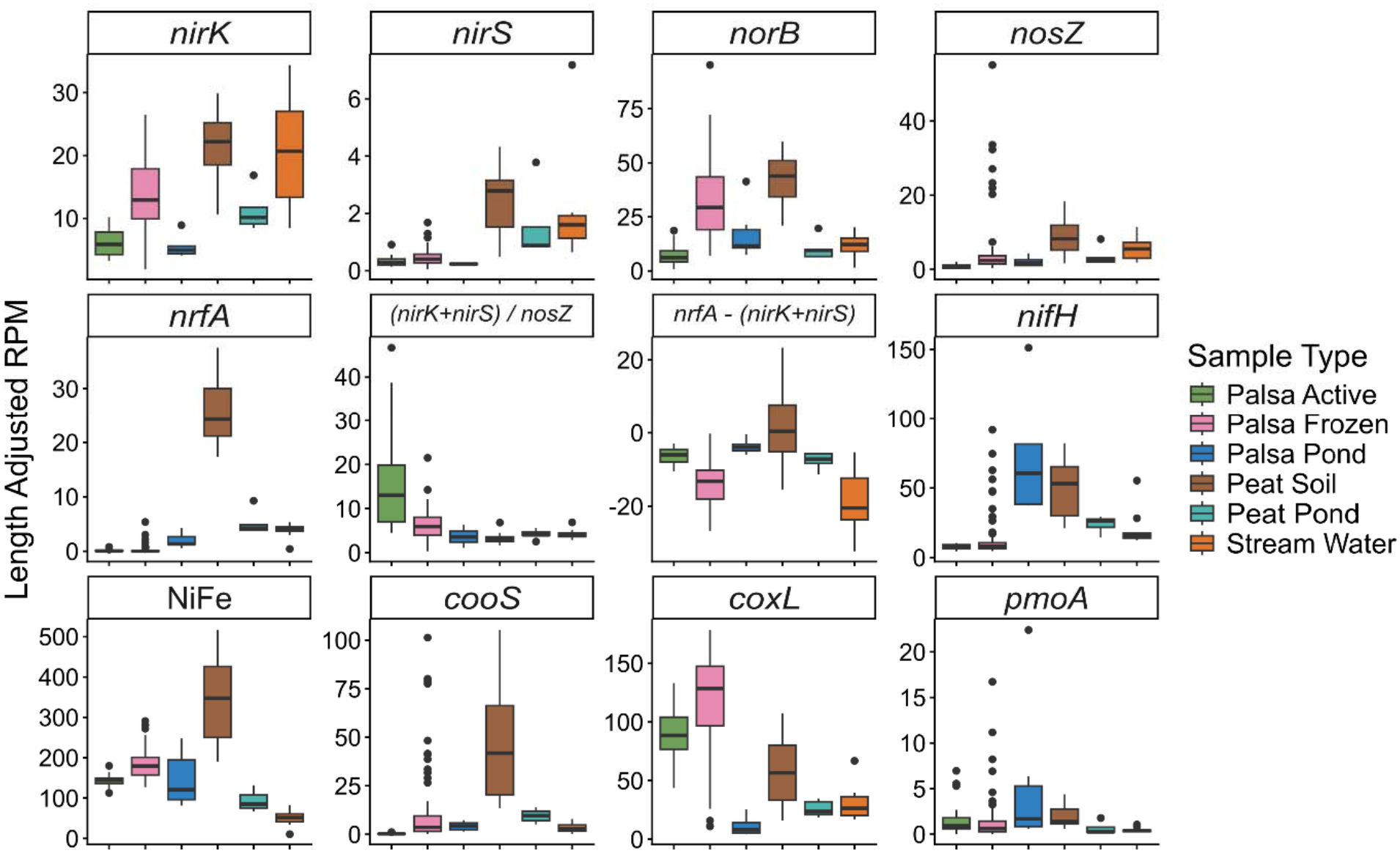
The abundance of selected metabolic marker genes in the sample types studied. The abundances of the metabolic marker genes are expressed as length adjusted reads per million (RPM) accounting for variations in gene length and sequencing depth. The marker gene ratios nirS+nirK/nosZ (ratio of potential of N_2_O production to N_2_O reduction) and nrfA - (nirK + nirS) (potential for nitrogen retention) were calculated from the length adjusted RPM values and are expressed as ratio or rate in the figure.

The gene *nrfA*, a marker gene for nitrate reduction to ammonium, had high abundance in the peat soils and the water samples when compared to the palsa soils or the water samples. Microbial community potential for nitrogen retention is indicated by the gene ratio of *nrfA*-(*nirK*+*nirS*) (Saghaï et al., 2023). The ratio was consistently negative across all palsa samples, both soil and ponds, as well as the stream water samples. However, in peat soils the mean ratio was positive (1.57 ±11.2 L-RPM). The gene *nifH*, a marker gene for nitrogen fixation, was highest in the peat soils and palsa ponds. There was increase in *nifH* in the palsa soil profile 26, with *nifH* reaching abundance of over 50 L-RPM in the deepest soil layers (>55cm, Figure S2).

#### Trace gas metabolism

Marker genes related to trace gas metabolism were widely detected across all of the sample types. [NiFe]-hydrogenase (*NiFe*), a marker gene for H_2_ utilization, had the highest value in the peat soils (340±110 L-RPM), followed by the palsa frozen soils (184 ±36 L-RPM), palsa ponds (146 ±69 L-RPM) and palsa active soils (142 ±16 L-RPM). *NiFe* had lower abundance in the peat ponds (93 ±26 L-RPM), and the stream waters (50 ±21 L-RPM).

Marker gene for aerobic CO oxidation (*coxL*) had the highest abundance in in the palsa frozen (116 ±42 L-RPM) and active soils (90 ±23 L-RPM), followed by the peat soils (57 ±30 L-RPM). On the other hand, the marker gene for anaerobic oxidation of CO (*cooS*) was highest in the peat soils (46 ±28 L-RPM), followed by palsa frozen soils (12 ±21 L-RPM).

The marker gene for methanogenesis (*mcrA*) had the highest average abundance in the peat soils (6.4 ±7.5 L-RPM), followed by palsa frozen soils (2.9 ±11.9 L-RPM) (Figure S2). The *mcrA* abundance increased in the soil profile 26 from <10 L-RPM to 40-60 L-RPM in the deepest layers >60 cm. Notably, the majority of palsa soil samples had no hits to *mcrA* (Table S2). In palsa active layer samples only 4 out of 20 had hits to *mcrA* and in frozen palsa soils 26 samples out of 72 had *mcrA* hits. Converserly, in peat soils, 14 out of 15 samples had hits to *mcrA*.

The aerobic methane oxidation marker gene *pmoA* was highest in the palsa ponds (5.6 ±8.5 L-RPM), followed by peat soils (1.9 ±1.1 L-RPM) and palsa active layer samples (1.9 ±2.1 L-RPM). The ratio of *pmoA*/*mcrA*, indicator of methane consumption to production was highest in the palsa frozen (27 ±52 L-RPM) and active soil samples (19 ±14 L-RPM), when taking into account only the samples for which the ratio could be calculated (Table S2). Out of the pond and stream samples, palsa ponds had the highest *pmoA*/*mcrA* ratio (6.0 ±5.9 L-RPM), followed by stream waters (3.2 ±3.3 L-RPM).

### Sample types hosted distinct microbiomes within the metagenome assembled genomes and only two genera present across all habitat types

Metagenome-assembled genomes (MAGs) were binned from metagenomes, yielding 1530 medium- and high-quality MAGs, of which 1345 were non-redundant (Table S3). The MAGs had an average completeness of 80% and contamination of 3%. The genome size average was 3 809 478bp (359 022-12 900 386bp) and CG content 0.60% (0.28-0.71%). 371 MAGs were high quality with completeness of ≥ 90% and contamination of <5% (Bowers et al., 2017). The mean genome size and GC content of the high-quality MAGs were on average 4 218 658bp and 0.58% respectively. The taxonomy of the MAGs spanned across 31 phyla, 71 classes and 392 genera. Genus level classification was missing from 3.9% of the MAGs (n=52, of which 20 were high-quality). The majority of the MAGs, 87% (n=1178), had no species level classification and no average nucleotide identity (ANI) value calculated by GTDBtk (Chaumeil et al., 2020). The most represented phyla were *Acidobacteriota* (n=327), *Actinomycetota* (n=293), *Pseudomodanota* (n=283), *Verrucomicrobiota* (n=80) and *Patescibacteriota* (n=57).

The most abundant phyla, with the highest abundance in TPM across all samples, were *Actinomycetota*, *Acidobacteriota*, *Pseudomonadota*, *Patescibacteriota* and *Chloroflexota* (Figure 4). *Patescibacteriota*, *Caldisericota* and *Halobacterota* were abundant in the deeper layers of the palsa soil profile 26, which was also in agreement with the read based results (Figure S1). The MAG based taxonomic composition was overall similar to the read based taxonomy. However, MAGs had higher relative abundance in phyla *Patescibacteriota, Babellota, Gemmatimonadota, Planktomycetota* and *Zixibacteria* compared to the read based results (Figure S3).

**Figure 4.**
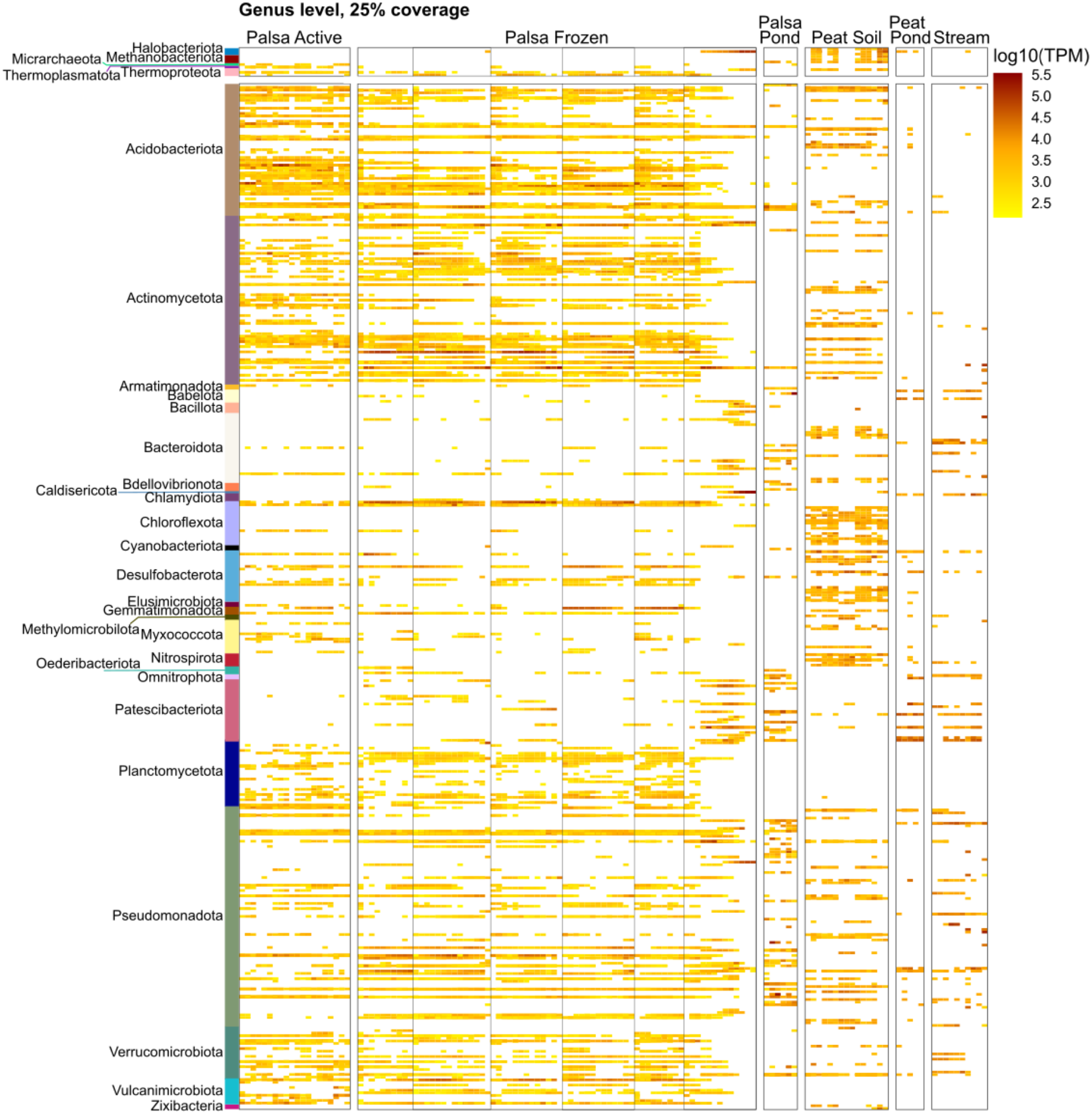
Heatmap of the genus level abundance normalized to transcripts per million based on the retrieved MAGs. The abundances are visualized based on the sample types of palsa active layer soils, palsa frozen layer soils, peat soils, palsa ponds, peat ponds and streams. Each row represents a genus, and each column a sample. The six replicate palsa soil profiles are separated by vertical lines in the palsa frozen group.

The community composition was distinct in palsa soils compared to the peat soils and water samples (Figure 5A), with a similar pattern to the read based results Figure 2). The archaeal phyla were divergent in palsa and peat soils, with *Thermoproteota* and *Thermoplasmatota* present in palsa soils and *Halobacterota*, *Thermoplasmatota* and *Methanobacterota* in peat soils. An exception in palsa soils was deepest layers of one palsa soil profile, where *Halobacterota* was the dominant archaea. *Desulfobacterota* and *Zixibacteria* were present in peat soils, but less abundant in the palsa soils. More precisely, only the class *Binatia* from *Desulfobacterota* was present in palsa soils, while in peat soils many classes of *Desulfobacterota* were detected. In the palsa soils *Planctomycetota* and *Vulcanimicrobiota* were present while mostly absent from the peat soils. *Chloroflexota* was abundant across all soil samples, with class *Dormibacteria* in palsa soils, and *Limnocylindria* and *Anaerolineae* in peat soils.

**Figure 5.**
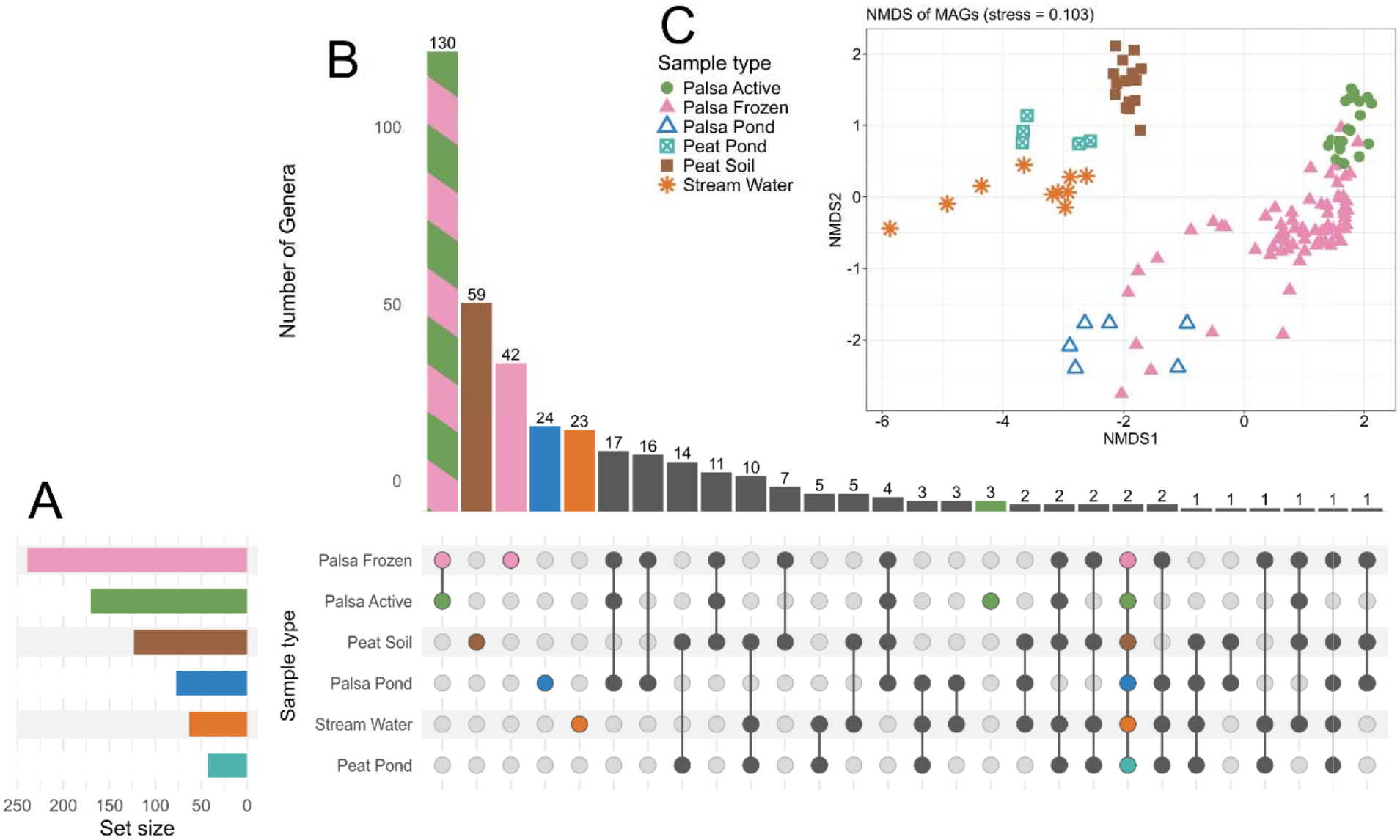
Number of genera per sample type (A) and the number of unique or shared genera across the sample types studied with the metagenome assembled genomes (MAGs) (B). Non-metric multidimensional scaling (NMDS) ordination of MAG abundances using Bray-Curtis dissimilarities based on TPM normalized abundance of the MAGs (C). MAGs with horizontal coverage of ≥25% were considered as present in this data.

The shared taxa among the sample types were explored with the presence of MAGs on genus level (Figure 5B). The only 2 genera shared across all sample types were *Rhizomicrobium* (phylum *Pseudomonadota*) and *UBA11358* (phylum *Verrucomicrobiota*). The palsa soils hosted the largest number of genera, which was expected due to the largest sample size. 130 genera were shared between palsa active and frozen soils, whereas active layer had three and frozen layers had 42 genera only detected in these sample types. Palsa ponds had 24 unique genera, 17 shared with both palsa soil types and 16 with frozen palsa soil. Peat soils had 59 unique genera, which was the highest number of unique genera out of all sample types. Peat soils and peat ponds shared 14 genera, whereas peat soils, peat ponds and streams shared 10 genera. Eleven genera were shared across the soil types, and three genera were present in peat ponds, palsa ponds and stream samples. No genera were shared only between palsa ponds and active layer soil or only between palsa soils, palsa ponds and stream waters (Figure 5B).

Interestingly, a symbiotic or parasitic lifestyle was relatively common among the detected MAGs across sample types. We detected phyla such as *Patescibacteriota, Babelota, Myxococcota, Omnitrophota* and *Bdellovibriota*. *Patescibacteriota* are endosymbionts (Bouderka et al., 2025) and were detected across all sample types. *Babelota* are intracellular parasites of protists (Weisse et al., 2025) and they were present mainly in the palsa ponds and streams. *Myxococcota* are predatory bacteria (Thiery & Kaimer, 2020) and were mainly detected in palsa and peat soils with classes *Myxococcia* and *Polyangia* being the most abundant. However, in palsa soil 26, class *UBA272* was the only *Myxococcia* detected. *Bdellovibrionota* was detected only in the frozen palsa soils and palsa ponds. Members of the class *Gorgyraeia* (*Omnitrophota*) were detected only in the stream samples, peat ponds and palsa ponds. *Gorgyraeia* are likely predatory bacteria (Seymour et al., 2023).

### Trace gas utilization is potential is across majority of the phyla and sample types

NiFe-hydrogenase was detected across wide range of classes (Figure 6C, Table S4), supporting the read based results (Figure 3) and also MAGs with NiFe were detected across all sample types. Similarly to the read-based analysis, potential for methanogenesis was detected in MAGs present in peat soils and one of the palsa soil profiles, by both marker genes (*mcrA*) and metabolic modules (Fig 6C-D).

**Figure 6.**
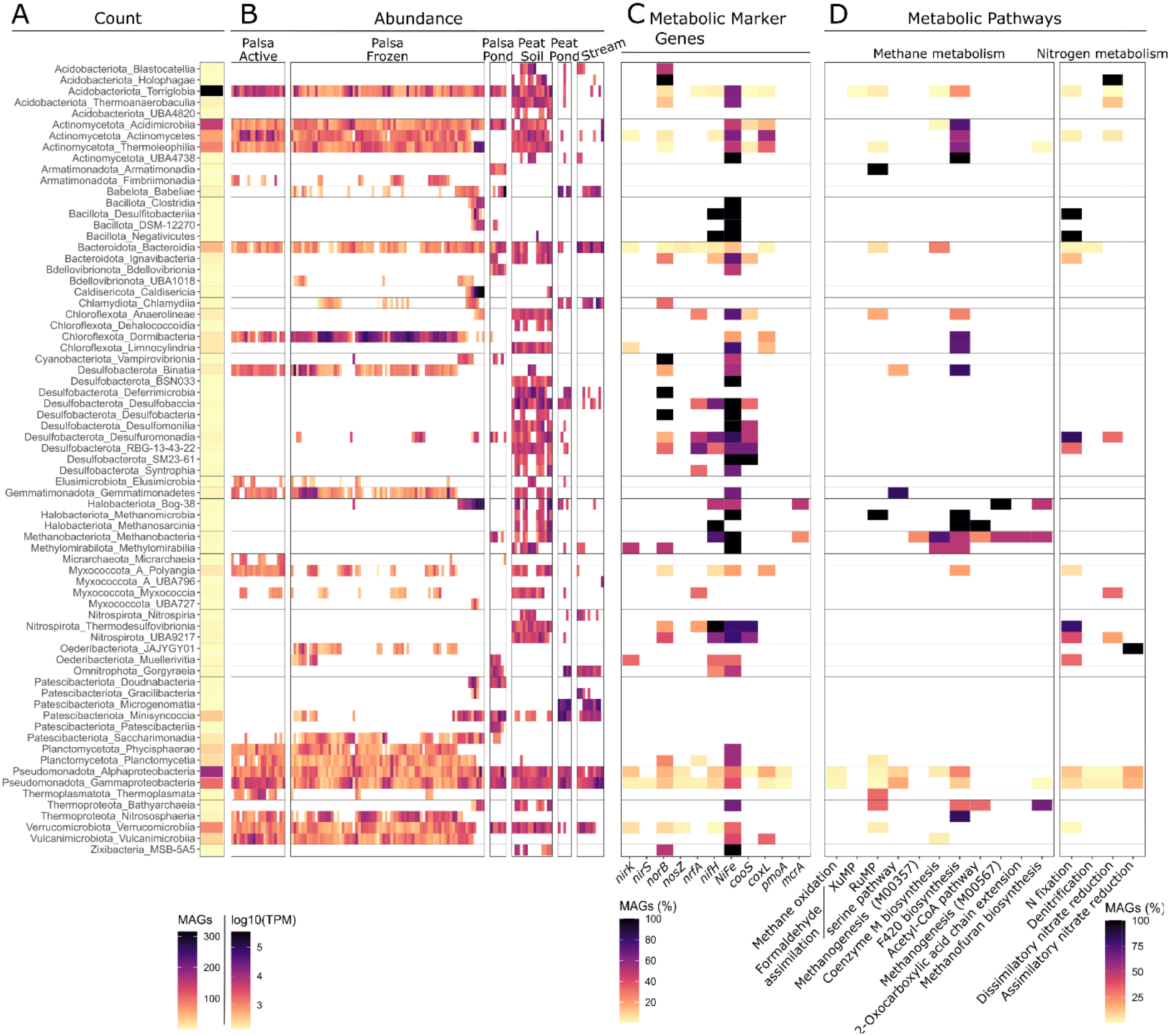
Number of dereplicated MAGs on each of the 71 bacterial and archaeal classes (A), their abundance across the sample types (B), and percentage of the MAGs per class with selected metabolic marker genes (C) and the KEGG metabolic pathways or modules of methane and nitrogen metabolism, which had pathway completeness of 75% (D).

Methanogenesis was detected in archaeal classes Methanobacteria and Bog-38. Marker genes for anaerobic CO oxidation (*cooS*) were detected in multiple classes of Desulfobacterota, Nitrospirota and Actinomycetota, as well as classes Terriglobia and Alphaproteobacterota. Many of these classes were mainly present in peat soil samples. However, classes with the aerobic CO oxidation marker gene *coxL*, were present in classes such as Dormibacterota and Vulcanimicrobiia, which were almost exclusively present in palsa soils. The presence of the nitrogen fixation marker gene *nifH* and complete nitrogen fixation pathway was detected across multiple classes, including the most numerous classes Terriglobia, Alphaproteobacteria and Gammaproteobacteria (Figure 6D, Table S5). While the full denitrification KEGG module was detected only in Pseudomonadota and Bacteroidota, the occurrence of separate dentification genes (such as *nirK* and *norB*) was more widespread in the studied sample types, spanning several different classes of MAGs. The potential for aerobic methanotrophy (*pmoA*) was detected in seven Pseudomonadota MAGs, of which 6 were present across palsa and peat soils and only one was detected in palsa ponds.

## Discussion

### Palsa soils host a unique microbiome compared to the peatlands and water samples

The aim of this work was to characterize microbial communities of a palsa peatland, including the transect of soil to ponds and streams. The most abundant taxa in our sampling sites were similar to previously described for permafrost peatland environments (Ernakovich et al., 2022; Hultman et al., 2015; Kirkwood et al., 2021; Waldrop et al., 2025; Waqas et al., 2026). Analysis of 107 peat and palsa soil metagenomes and 21 aquatic metagenomes yielded 1 345 metagenome assembled genomes, of which majority could not be assigned a species level taxonomy or calculated an ANI value. These genomes originating from metagenomes will expand the catalog of bacterial and archaeal genomes from northern peatland and palsa, including some ubiquitous phyla such as *Vulcanimicrobiota, Omnitrophota, Elusimicrobiota, Oederibacteriota* (prev. *FCPU426*), *Patescibacteriota*, *Zixibacteriota* and *Babelota*. There were also multiple host dependent bacterial phyla detected, similarly to Antarctic soils (Ortiz et al., 2021). These included *Bdellovibrionota*, *Babellota* (prev. *Dependentiae*), *Chlamydiota* and *Patescibacteriota*, all of which have predatory, intracellular or symbiotic lifestyles.

The majority of the genera in MAGs were only detected in one of the sample types, indicating unique microbiomes even between the soils and their adjacent ponds and thaw ponds. Palsa soils were distinct from the peat soils and water samples in their microbial community composition. Leroy et al. (2026) detected decreasing similarity between palsa soils and the forming thaw ponds as the ponds matured. In our sites, around two thirds of the genera detected in the palsa thaw ponds were also present in palsa soils, and one third was unique to the thaw ponds, whereas less connectivity was detected between the palsa ponds and streams. In peat soils there was more of an overlap in taxonomy between soils, ponds and the streams draining the site. This was likely due to a higher degree of hydrological connectivity in the peatland compared to the palsa soils.

### Metabolic marker gene abundance suggested loss of nitrogen and increased potential of N_2_O emissions from palsa soils

The gene ratio (*nirS*+*nirK)*/*nosZ* ratio, indicator of N_2_O production over N_2_O consumption, indicated potential for N_2_O emissions across samples as the ratio was consistently higher than one across samples. The ratio in palsa frozen layers, peat and water sites was of similar range as in Yedoma permafrost (Marushchak et al., 2021) and northern peatlands (Välikangas et al., 2026). However, the active layer of palsa had a ratio of 3-4 times higher than the Yedoma or peatland soils, suggesting higher potential for incomplete denitrification and higher N_2_O emissions. Based on the marker gene analysis it is plausible that N_2_O will be emitted from the palsa soils as they thaw, as has been shown previously in this palsa site (Bhattarai et al., 2022) and in palsa soils elsewhere (Burnett et al., 2026). However, multiple MAGs encoded either full denitrification module or some of the related marker genes. Denitrification is largely based on cross feeding between microbes with parts of the metabolic pathway present (Pessi et al., 2022), and thus complete denitrification is also likely also occurring across sample types.

### H_2_ and CO utilization was prevalent in palsa and peat soils, while methane metabolism was observed in peat soils

Utilization of H_2_ and CO was detected across all soil types and all depths. Trace gas utilization is likely vital for microbial communities in palsa and peat environments. The ability to oxidize H_2_ and CO is detected widely in the phyla dominating the soils, including *Acidobacteriota, Actinobacteriota, Bacteroidota, Chloroflexota, Firmicutes, Gemmatimonadota, Planctomycetota*, *Proteobacteria* and *Verrucomicrobiota* (Bay et al., 2021). In this study, the ability to oxidize H_2_ was also detected widely across the MAGs in these phyla, but also in three archaeal phyla and across 17 bacterial phyla. In addition, the oxidation of CO was also detected in a wide range of classes across ecosystems, supporting the finding that trace gas utilization is highly prevalent in this palsa ecosystem. The ability to utilize both organic and inorganic energy sources is beneficial to microbes, as they can flexibly utilize inorganic energy sources if organic sources are depleted or limited (Bay et al., 2021). While high in organic matter and nutrients, these resources might be unavailable in palsa soils due to the frozen state of the soil. Apart from resource availability, the high potential for trace gas utilization across the frozen soils could be due to dormancy (Greening & Grinter, 2022).

Potential for aerobic methane oxidation was detected in palsa ponds and some of the palsa and peat soils, in which also the MAGs with *pmoA* gene mainly from class *Rhizobiales* (*Pseudomonadota*) were detected. Potential for methanogenesis was not detected in the majority of the palsa soil layers, which is in agreement with the low abundance or lack of methanogenic archaea according to both read-based analysis and MAGs in these sample types. The marker gene for metanogenesis (*mcrA)* was abundant only in one palsa soil profile 26, which also had increased abundance of *Halobacterota*. As expected, the highest potential for methanogenesis was detected in waterlogged peat soils. Similarly, only the peat soils indicated potential for higher methane production than consumption.

While considering the lack of methanogenesis and the ratio between methanogenesis and methanotrophy in our water samples, these samples were collected from the water column and not from the sediment, which can be the main source of methane in palsa and peat ponds (Knutson et al., 2025; Leroy et al., 2026). Moreover, the sites were sampled during late snowmelt, when the methanogenic community may not be fully formed, as methanogens are most abundant in the summer season (Vigneron et al., 2019).

### One palsa soil profile was thawing, leading to changes in community and metabolic potential

One notable pattern occurring across the analysis was the palsa soil profile 26, which differed from rest of the palsa soils in pH, read based taxonomy, MAG abundance, and metabolic marker gene counts. The deepest soils of palsa profile 26 had community composition more similar to the palsa ponds and peat soils than to the other palsa soil profiles. There was higher also abundance of genus *Methanoflorens* (*Halobacterota*) in this one palsa soil profile compared to the other palsa profiles. *Methyloflorens* is a known hydrogenotrophic methanogen, which has been previously described in degrading permafrost peatland (Ellenbogen et al., 2024; Hultman et al., 2015). This coincided with the soil depths where pH increased from 4 to 5 and higher counts of metabolic marker genes *mcrA* and *nifH*. Increase in pH in palsa soils has been linked to increase in CH_4_ emissions (Niemi et al., 2026), supporting our findings. In addition, these soil layers had higher abundances of *Patescibacteriota*, *Caldisericota* and *Thermoproteota*.

Based on our results, we conclude that this palsa soil was likely to thaw for above and below, leading to low oxygen conditions and increased connection with the surrounding thaw ponds and mineral ground water. This palsa soil profile is highly likely to have been anaerobic, considering the results supporting methanogenesis and nitrogen fixation. N_2_O emissions in palsa soil are expected to decrease as the soil collapses and gets into contact with mineral groundwater and increases the soil pH (Takatsu et al., 2022). However, this pH increase was not detected in the other palsa soils profiles, except the soil profile 26.

## Conclusions

Palsa soils and ponds have a unique microbiome on both sequence read and MAG level when compared to nearby peatland soil, peat ponds and stream samples. The soil and adjacent pond shared genera as expected in both palsa and peatland sites. However, there were no taxa shared only between palsa soils, ponds and streams. The peat soil, peat ponds and streams did share genera, indicating better connectivity between these sample types. Multiple MAGs with potentially symbiotic or parasitic lifestyles were detected across all sample types, indicating the potential importance of symbiotic lifestyles in the peat and palsa ecosystem. The potential for trace gas metabolism, especially utilization of hydrogen, was detected widely across the sample types and metagenome assembled genomes. Based on metagenomes, the palsa soils had potential to be a source of N_2_O, whereas peatlands were a possible source of methane and a sink for nitrogen.

## Author contributions

Nea Tuomela (Conceptualization, Funding acquisition, Field work, Data curation, Formal analysis, Investigation, Visualization, Writing—original draft, Writing—review & editing), David N. Thomas (Conceptualization, Project administration, Field work, Investigation, Resources, Supervision, Writing—review & editing), Hermanni Kaartokallio (Conceptualization, Supervision, Investigation, Resources, Writing— review & editing), Tommi Välikangas (Data curation, Writing – review & editing), and Jenni Hultman (Conceptualization, Supervision, Investigation, Resources, Writing— review & editing).

## Data availability

The data is under embargo until publication. Sequencing data will be made available in ENA projects PRJEB121465 and PRJEB85624. Metagenome assembled genomes will be made accessible in Figshare https://doi.org/10.6084/m9.figshare.34898898.

## Acknowledgements

We thank Kevo Subarctic Research Station (University of Turku), Samu Elovaara and Janina Kariluoto for assistance with the field campaign. We also thank Igor Pessi and Sirja Viitamäki for assistance with computational analysis. We thank CSC—IT Center for Science for the computational resources. We acknowledge DNA Sequencing and Genomics Laboratory (supported by HiLIFE and Biocenter Finland funding), Institute of Biotechnology, University of Helsinki for sequencing.

