## Supplementary Methods and Figures for "Northern palsa and peat ecosystem microbiomes are potentially supported by hydrogen and carbon monoxide metabolism"

Authors & affiliations:

Nea Tuomela^1*^, David N. Thomas^1^, Hermanni Kaartokallio^2^, Tommi Välikangas^3^ and Jenni Hultman^3^

^1^ Department of Environmental Sciences, Faculty of Biological and Environmental Sciences, University of Helsinki, Helsinki, Finland

^2^ Marine and Freshwater Unit, Finnish Environment Institute, Helsinki, Finland

^3^ Soil Ecosystems, Natural Resources Institute Finland, Helsinki, Finland

*Corresponding author Nea Tuomela, Department of Environmental Sciences, Faculty of Biological and Environmental Sciences University of Helsinki, Viikinkaari 1, 00790 Helsinki,

### Supplementary Methods

**Read based marker gene analysis**

To investigate the metabolic functional potential in the data, the metagenomic reads were annotated using the curated 51 metabolic marker gene database from Greening Lab (Leung and Greening 2021, Lappan et al. 2023) similar to (Välikangas *et al.* 2026) . Briefly, the paired-end reads were independently aligned for forward (R1) and reverse (R2) reads against the compiled metabolic marker gene database with DIAMOND (Buchfink, Xie, and Huson 2014) blastx v. 2.1.16 (--query-cover 80 --max-target-seqs 1 --outfmt 6). The alignment results from R1 and R2 were subsequently filtered independently using marker gene specific minimum percentage identity thresholds set to 80% (psaA), 75% (hbsT), 70% (psbA, isoA, atpA, ygfK, aro), 60% (coxL, mmoA, amoA, nxrA, rbcL, nuoF, fefe hydrogenases, nife Group 4 hydrogenases) and 50% (all other genes), following (Lappan et al. 2023).

The filtered R1 and R2 alignments were reconciled at the paired-fragment level so that each sequenced fragment contributed at most to one marker gene assignment. In short:

1. For read pairs where only one mate (R1 or R2) produced a valid marker gene hit, the corresponding alignment was retained.
2. If both mates aligned to the same marker gene, the better supported alignment was retained according sequentially to higher bitscore, higher percentage identity when bitscores were tied, and higher query coverage when both bitscore and percentage identity were tied. If all three measures were tied, the R1 alignment was retained.
3. If mates aligned to different marker genes, pairs for which both alignments had bitscores <50 were discarded. Otherwise, if the absolute bitscore difference was ≥15, the alignment with the higher bitscore was retained. Discordant pairs with an absolute bitscore difference <15 were considered ambiguous and discarded.

With this approach, each retained paired-end fragment contributed a maximum count of one, preventing double counting when both mates produced valid alignments.

The retained marker gene alignment hits were further adjusted to account for variation in gene length for the marker gene database proteins and differences in sequencing depth between the samples. First, the amino-acid length of the matched marker gene database protein was converted to an approximate coding-sequence length by multiplying the protein length by three and expressing the resulting length in kilobases. Reads per kilobase (RPK) were then calculated by dividing the alignment hit count by this approximate coding-sequence length in kilobases. Next, the total number of sequenced paired-end fragments in each sample was used to calculate a library-size-per-million scaling factor as the total number of sequenced fragments / 10^6^. A length- and sequencing-depth normalized abundance measure, length adjusted reads per million (length adjusted RPM), was calculated by dividing the RPK values by this library-size-per-million scaling factor. The resulting length adjusted RPM values were then aggregated by marker gene.

The acquired length adjusted RPM differs from the commonly used transcripts per million (TPM) metric for RNA-Sequencing primarily in its scaling denominator. The used length adjusted RPM is scaled by the total number of sequenced fragments in the sample / 10^6^. In contrast, if TPM were calculated within the quantified marker gene dataset, the corresponding used scaling factor would be the sum of RPK values across all quantified marker genes in that sample / 10^6^. Consequently, the length adjusted RPM values are not constrained to sum to a constant value across the marker gene set, and changes in the abundance of one marker gene do not directly rescale the values of the other marker genes through a marker gene set specific denominator. This avoids the additional compositional closure that would result from calculating TPM within the small marker gene subset, although the resulting length adjusted RPM values remain relative sequencing abundances rather than absolute gene abundances.

On the other hand, TPM values calculated directly within the marker gene subset would sum to 10^6^ in each sample and describe the relative contribution of each marker gene to the total length normalized abundance represented by the quantified marker gene subset. Thus, length adjusted RPM can be interpreted as the length normalized abundance of each marker gene per million sequenced fragments (sequencing depth), whereas TPM calculated within the 51 marker gene set would describe each marker gene's share of the total length normalized abundance of that marker gene set.

### Supplementary Figures

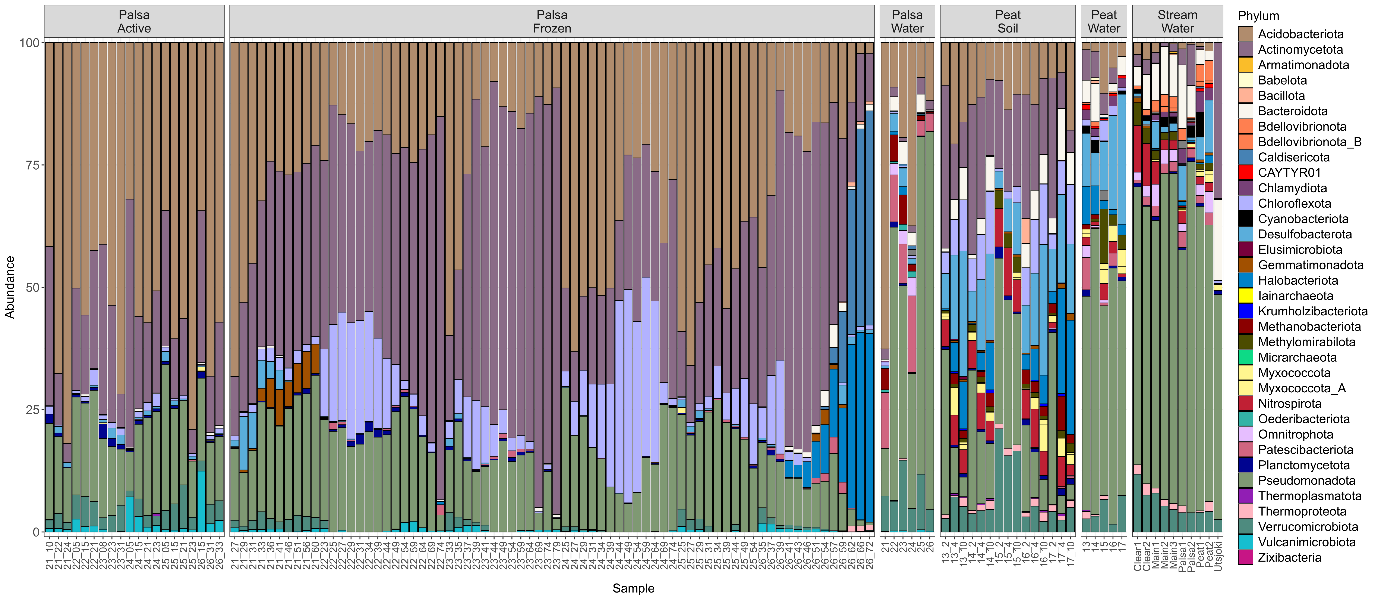

Figure S1. Microbial community composition on phylum level from metagenomes of palsa and peat soils, ponds and stream water samples. X-axis has sample name for all, and for soil samples, the sample id is followed by sampling depth ( cm).

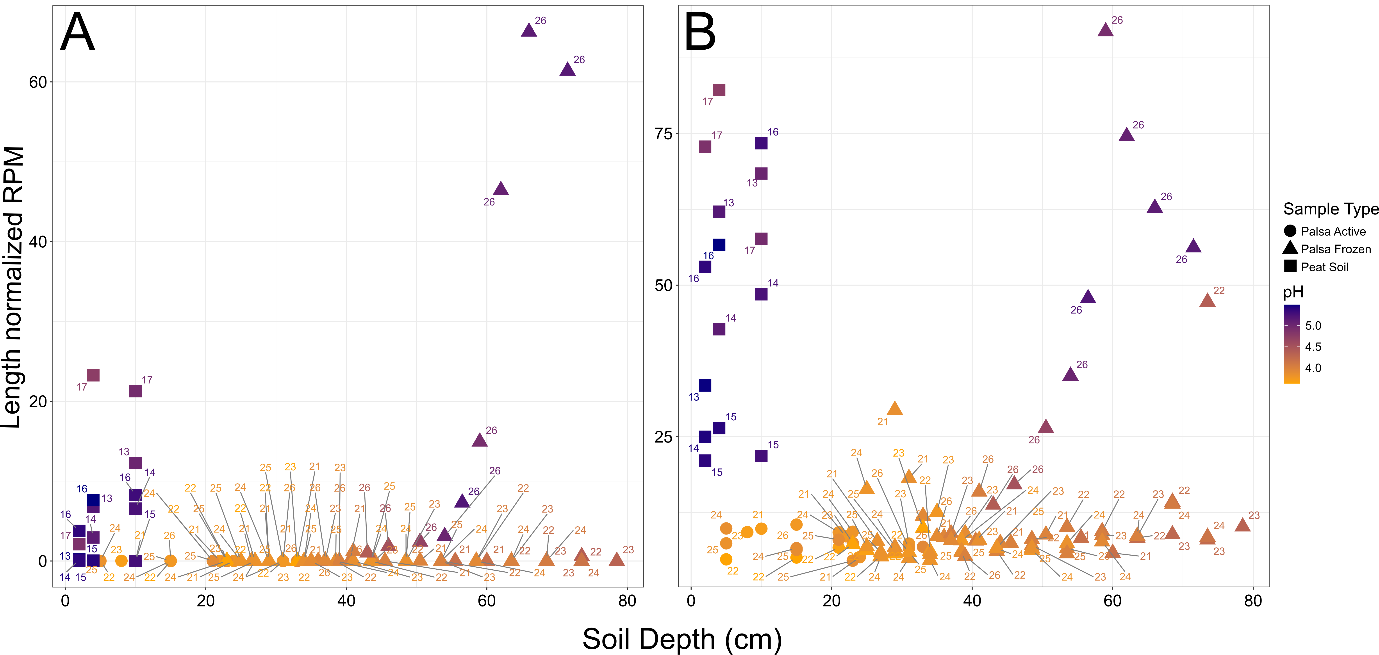

Figure S2. The abundance of metabolic marker genes for methanogenesis *mcrA* (A) and nitrogen fixation *nifH* (B) in peat and palsa soils samples by depth. The marker gene counts are normalized to length adjusted reads per million (RPM). Labelling indicates sample id, shape refers to sample type and colour for pH value.

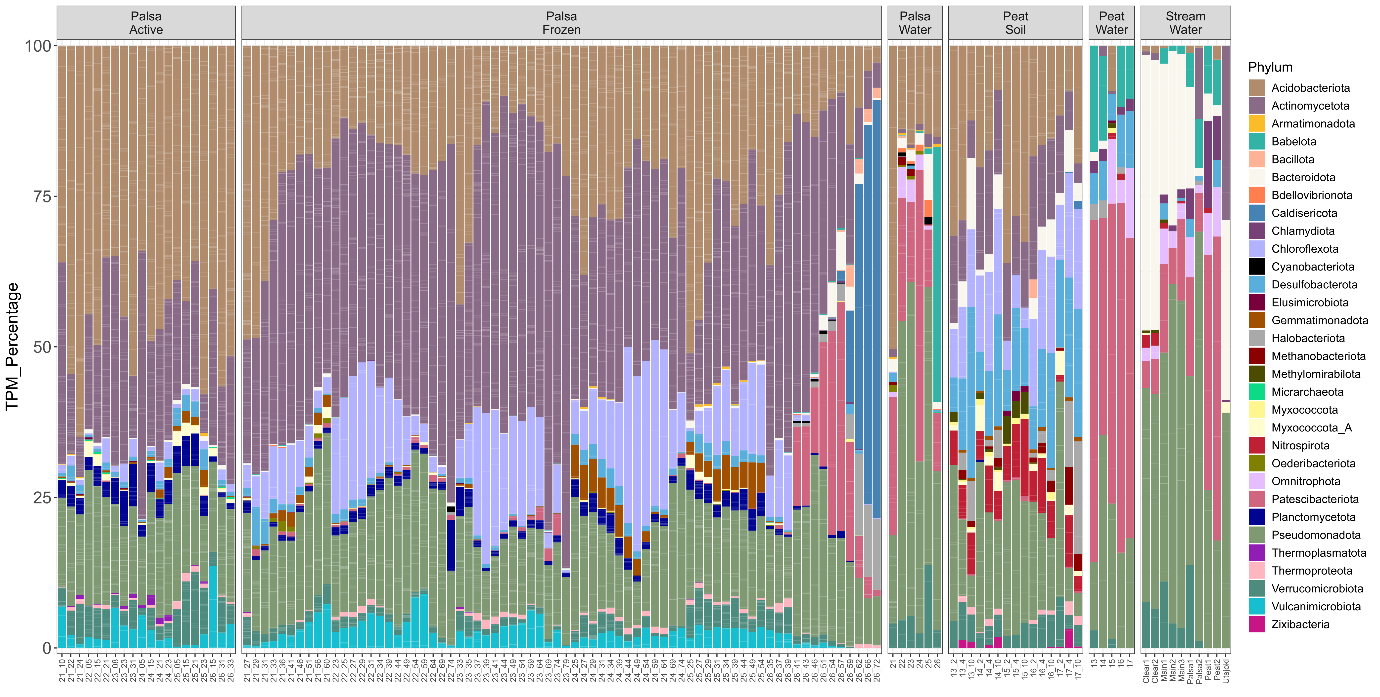

Figure S3. Microbial community composition in phylum level based on the abundance of metagenome assembled genomes. X-axis has sample name for all, and for soil samples, the sample id is followed by sampling depth (in cm).
